# Leveraging medical Twitter to build a visual–language foundation model for pathology AI

**DOI:** 10.1101/2023.03.29.534834

**Authors:** Zhi Huang, Federico Bianchi, Mert Yuksekgonul, Thomas Montine, James Zou

## Abstract

The lack of annotated publicly available medical images is a major barrier for innovations. At the same time, many de-identified images and much knowledge are shared by clinicians on public forums such as medical Twitter. Here we harness these crowd platforms to curate OpenPath, a large dataset of 208,414 pathology images paired with natural language descriptions. This is the largest public dataset for pathology images annotated with natural text. We demonstrate the value of this resource by developing PLIP, a multimodal AI with both image and text understanding, which is trained on OpenPath. PLIP achieves state-of-the-art zero-shot and transfer learning performances for classifying new pathology images across diverse tasks. Moreover, PLIP enables users to retrieve similar cases by either image or natural language search, greatly facilitating knowledge sharing. Our approach demonstrates that publicly shared medical information is a tremendous resource that can be harnessed to advance biomedical AI.

## Introduction

Recent advances in artificial intelligence (AI) algorithms in computational pathology can help to distinguish cell or tissue types, generate diagnoses, and retrieve relevant images from routinely stained hematoxylin and eosin (H&E) images [1–5]. However, computational progress is heavily bottlenecked by the insufficient number of well-annotated pathology images, as the scale of publicly available annotated pathology datasets falls below the common standards in other AI domains [6]. This limitation is especially problematic in pathology where there are more than 8,000 diseases [7] to consider, and their pathologic classification is constantly evolving as knowledge of the molecular and cellular bases of disease advances [8].

At the same time, many de-identified pathology images are shared on the Internet, especially on social media, where clinicians discuss de-identified medical images with their colleagues [9–12]. These public data and discussions hold significant value to the pathology community, enabling knowledge-sharing and serving educational purposes. In particular, there exists a comprehensive collection of pathology subspecialty-specific hashtags in the Twitter community, which were user-generated and systematically arranged by the 2016 United States and Canadian Academy for Pathology (USCAP) meeting [13,14]. This set of data includes both common and rare pathology cases, which were routinely stained with H&E dyes, and sometimes by immunohistochemistry (IHC). These public discourses represent an underutilized source of data and knowledge for medical AI [15–17].

In this paper, we used the popular pathology Twitter hashtags to curate 243,375 public pathology images. We expanded this collection to include pathology data from other sites on the Internet (collected from LAION [18]), followed by strict data quality filtering, finally created a collection of 208,414 pathology image–text pairs called OpenPath. To our knowledge, OpenPath is so far the largest publicly available pathology image collection that is annotated with text descriptions. We then leverage this large-scale structured set of pathology image–text pairs to develop a versatile image-and-language AI foundation model for pathology.

This study first presents a complete description of the collected OpenPath dataset and proposes PLIP (Pathology Language–Image Pre-training) model, which was trained on paired images and captions from OpenPath via contrastive learning. Next, a comprehensive evaluation of our proposed model is conducted to assess its ability to adapt to new captions through zero-shot learning [19]. Different from other pathology image classification models which were trained from a fixed set of categorical labels, such as binary classification [20,21], semantic segmentation for tumors [22], identifying a fixed number of tissue types [23,24], binary or multi-class differential diagnosis [25,26], the proposed zero-shot learning from PLIP can perform classification without having been specifically trained to classify them, making it useful as a versatile system for the potential emergence of new disease subtypes. Moreover, PLIP can function as a general purpose image encoder to capture a better image representation for pathology, making it possible to train and classify new sets of data via linear probing, leading to improved performances across diverse tissue types and learning tasks. Finally, PLIP enables a flexible search engine for pathology images. We conducted a systematic evaluation of image retrieval [4,5] to demonstrate its ability to retrieve relevant pathology images by text or image inputs, which holds significant knowledge-sharing potential.

## Results

### Creating OpenPath from Twitter and other public sources

The United States and Canadian Academy for Pathology (USCAP) and the Pathology Hashtag Ontology projects (https://www.symplur.com/healthcare-hashtags/ontology/pathology/) recommends 32 Twitter pathology subspecialty-specific hashtags [13,14]. We used these 32 hashtags to retrieve relevant tweets from 2006-03-21 (the date of the first Twitter post) to 2022-11-15 (**Figure 1a**) to establish so far the largest public pathology dataset with natural language descriptions for each image: OpenPath. The detailed definition of each hashtag is presented in **Extended Table 1**. We followed the usage policy and guidelines from Twitter and other entities in retrieving the data. To ensure data quality, OpenPath followed rigorous protocols for cohort inclusion and exclusion, including the removal of retweets, sensitive tweets, and non-pathology images, as well as additional text cleaning (**Figure 1a**, **Extended Figure 1** and **Online Methods**). The final OpenPath dataset (**Figure 1b**) consists of (1) *Tweets*: 116,504 image–text pairs from Twitter posts (tweets) across 32 pathology subspecialty-specific hashtags (**Figure 1c**); (2) *Replies*: 59,869 image–text pairs from the associated replies that received the highest number of likes in the tweet, if applicable (**Figure 1c**); and (3) *PathLAION*: 32,041 additional image–text pairs scraped from the Internet and the LAION dataset. The detailed dataset extraction and description are further elaborated in the **Online Methods** section, and the complete dataset inclusion–exclusion procedure is demonstrated in **Extended Figure 1**.

**Figure 1.**
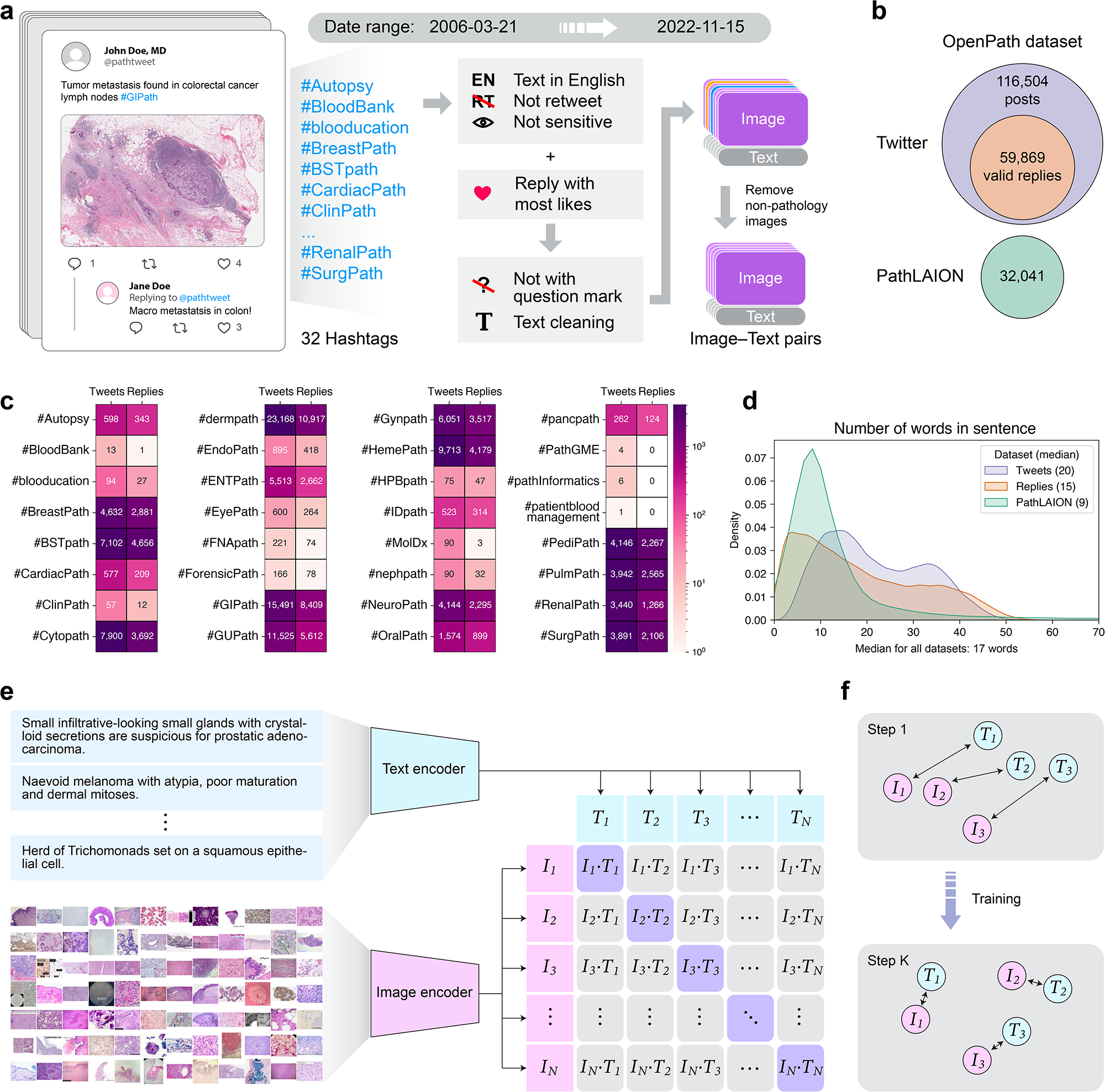
Overview of this study. **(a)** Flowchart of data acquisition from medical Twitter. **(b)** Overview of the OpenPath dataset. **(c)** Total number of available image–text pairs from tweets and replies within each Twitter hashtag (sorted in alphabetical order). Replies are those which received the highest number of likes in Twitter posts, if applicable. **(d)** Density plot of the number of words per sentence in the OpenPath dataset. **(e)** The process of training the PLIP model with paired image–text dataset via contrastive learning. **(f)** Graphical demonstration of the contrastive learning training process.

For quality assurance, an additional manual inspection was conducted by randomly sampling 1,000 images from the OpenPath dataset (see **Extended Table 2**). This evaluation showed that 98.5% of the subsampled images were pathology images and 94.5% of the sampled images were high-quality regions of interest from whole slide images, indicating that the OpenPath contains data of reasonably good quality. Furthermore, the captions in OpenPath had a median number of 17 words (**Figure 1d**, **Extended Table 3**) and contained high-quality descriptions of the medical condition in the image. In the next sections, we describe how we leverage OpenPath to develop the state-of-the-art foundation visual–language model for pathology.

### Training a visual–language AI using OpenPath

While visual–language models have been successful in a myriad of natural image tasks, their applicability to domain-specific tasks, such as medical tasks, have been limited. With detailed natural language annotations, OpenPath can be used to train a visual–language foundation model that can perform well on many medical AI applications. Unlike other supervised learning and segmentation pathology models that were trained solely on categorical labels, texts in natural language are enriched with semantic and interrelated knowledge, which can further enhance the understanding of the images and facilitate multiple downstream applications. In this study, we fine-tuned a pre-trained CLIP model [27] on OpenPath using contrastive learning. To accomplish this, a pathology image pre-processing pipeline was integrated, including image downsampling, random cropping, and data augmentations (**Online Methods**). During the training phase, the PLIP model generates two embedding vectors from both the text and image encoders (**Figure 1e**). These vectors were then optimized to be similar for each of the paired image and text vectors and dissimilar for non-paired images and texts via contrastive learning (**Figure 1f**, **Online Methods**).

Leveraging the advantages of contrastive learning and the detailed text descriptions from OpenPath, PLIP is able to handle multiple types of inferences across a broad spectrum of medical applications. This capability, which does not require explicit training, sets PLIP apart from other supervised counterparts. For instance, when presented with an image and multiple disease descriptions, PLIP can determine which candidate description best describes that given image. Additionally, given a disease or tissue description, PLIP can retrieve the most relevant images that best align with the input description. In contrast, supervised models are restricted to making predictions based on the categorical labels they were trained on, limiting their ability to perform tasks beyond their training scopes. These unique attributes in PLIP significantly streamline the efforts to implement or deploy AI for digital pathology under complex situations for various downstream tasks. In subsequent sections, we demonstrate how PLIP can be used to perform zero-shot learning, and how it can be used as a pretrained model for transfer learning, as well as its ability to search image databases using text and image queries.

### PLIP can perform zero-shot classification on new data

Benefiting from its understanding of the text, the proposed PLIP model can classify and recognize new labels for unseen data. This capability, commonly referred to as zero-shot learning [19], enables learning new classes at scale without the need for re-training. In zero-shot learning, the classification result is based on identifying the candidate texts with the highest representational similarity to the input image (**Figure 2a**).

**Figure 2.**
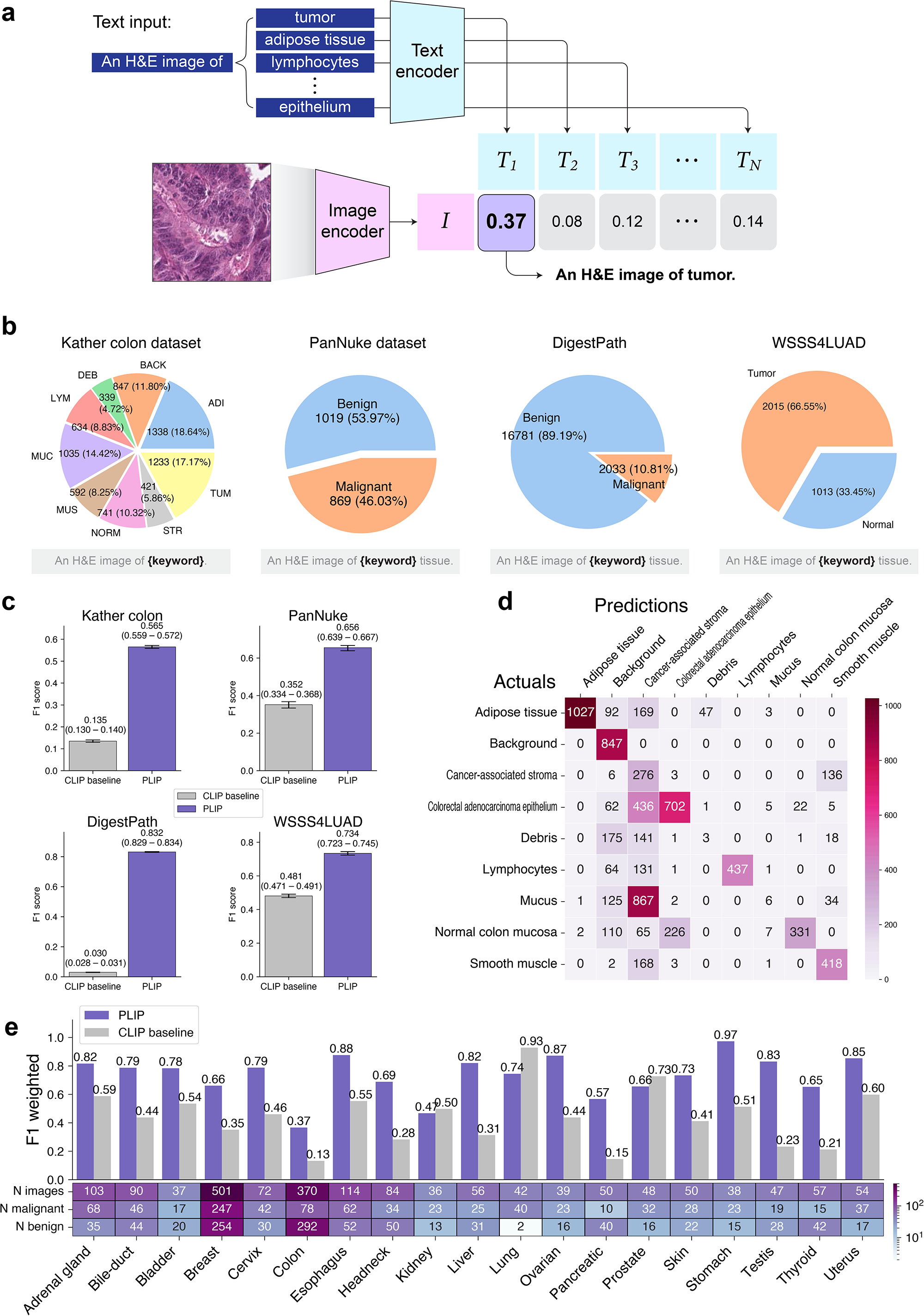
PLIP predicts new classes via zero-shot transfer learning. **(a)** Graphical illustration of zero-shot classification. The classification output is determined by selecting the candidate text with the highest cosine similarity to the input image. **(b)** Four external validation datasets: Kather colon dataset with 9 tissue types, PanNuke dataset (benign and malignant tissues), DigestPath dataset (benign and malignant tissues), and WSSS4LUAD dataset (tumor and normal tissues). In the Kather colon dataset, ADI: Adipose tissue, BACK: background, DEB: debris, LYM: lymphocytes, MUC: mucus, MUS: smooth muscle, NORM: normal colon mucosa, STR: cancer-associated stroma, TUM: colorectal adenocarcinoma epithelium. **(c)** Zero-shot performances with weighted F1 scores across four datasets. Note that the performances in the Kather colon dataset are based on a 9-class zero-shot learning evaluation, while the performances for other datasets are on binary zero-shot learning evaluation. **(d)** Confusion matrix of Kather colon dataset. The actual and predicted labels are displayed in rows and in columns, respectively. **(e)** Zero-shot evaluation of the PanNuke dataset within each organ type.

In this study, we conducted a systematic evaluation of PLIP’s zero-shot capability across four external validation datasets: (i) the Kather colon dataset [24] with 9 different tissue types; (ii) the PanNuke dataset [28] with two tissue types (benign and malignant); (iii) DigestPath dataset [29] with two tissue types (benign and malignant); and (iv) WSSS4LUAD dataset [30] with two tissue types (tumor and normal) (**Figure 2b**, **Extended Figure 2**). To evaluate the PLIP model on those datasets, labels were converted to sentences. For example, “tumor” was converted to “An H&E image of tumor”. As a natural comparison on the zero-shot task, we compared PLIP with the original CLIP model, which has been frequently used for other medical image tasks [31–33] and has already been trained from other medical images. Our analysis showed that PLIP consistently outperformed the baseline CLIP model and the results from predicting the majority class (or Majority) (**Figure 2c**). For instance, PLIP achieved F1 = 0.565 (95% CI = 0.559 – 0.572) on the Kather colon dataset (9 classes), which is 4.2x and 9.6x higher than CLIP (0.135) and Majority (0.059), respectively. In the PanNuke dataset (benign vs. malignant), PLIP achieved F1 = 0.656 (95% CI = 0.639 – 0.667), which is 1.9x and 1.7x higher than CLIP (0.352) and Majority (0.379), respectively. In the DigestPath dataset (benign vs. malignant), PLIP achieved F1 = 0.832 (95% CI = 0.829 – 0.834), which is 27.7x higher than CLIP (0.030) and similar to Majority (0.841). In WSSS4LUAD (tumor vs. normal), PLIP achieved F1 = 0.734 (95% CI = 0.723 – 0.745), which is 1.5x and 1.4x higher than CLIP (0.481) and Majority (0.532), respectively.

In addition, the confusion matrix with ground truth and predicted annotations for the Kather colon dataset is presented in **Figure 2d**. Compared to other models’ performance (**Extended Figure 3**), this result suggested that PLIP achieved a reasonable zero-shot learning capability on this sophisticated task. While our proposed PLIP model demonstrated the ability to accurately distinguish several key tissue types, including adipose tissue, background, colorectal carcinoma epithelium, and lymphocytes, it may face challenges when dealing with other tissue types, such as mucus and debris. This observation is likely due to mucus, debris, and smooth muscle sometimes being considered cancer-associated stroma.

Moreover, the binary zero-shot outcomes from the PanNuke dataset drew our attention and prompted us to conduct a more in-depth investigation into the zero-shot performances to determine benign vs. malignant among samples within each of 19 different organs in PanNuke dataset. Compared to baseline CLIP in **Figure 2e**, we found PLIP achieved superior weighted F1 score on 16 out of 19 organs, namely: adrenal gland (0.82), bile-duct (0.79), bladder (0.78), breast (0.66), cervix (0.79), colon (0.37), esophagus (0.88), head-neck (0.69), liver (0.82), ovarian (0.87), pancreatic (0.57), skin (0.73), stomach (0.97), testis (0.83), thyroid (0.65), and uterus (0.85). Among them, 7 subspecialties (adrenal gland, esophagus, liver, ovarian, stomach, testis, and uterus) achieved reasonably high F1 scores (>0.8), whereas the baseline CLIP performed only at F1 = 0.3 ∼ 0.6. However, imbalanced classes were observed among samples from some organs, such as kidney, lung, and prostate, and this may have contributed to inferior performances by PLIP.

### PLIP provides better image representations for training models

To gain a deeper understanding of the capabilities of the PLIP image encoder, we used four different testing datasets (Kather colon [24], PanNuke [28], DigestPath [29], and WSSS4LUAD [30]) to evaluate the ability of image representation. Image embeddings were first calculated by the PLIP image encoder and then applied dimensionality reduction via UMAP (**Figure 3a–d**). Despite that these testing pathology images were previously unseen by the PLIP model and might have a different data distribution from what PLIP was trained on, we found PLIP can still effectively distinguish between various tissue subtypes in the Kather colon dataset (**Figure 3a**). Compared to other baseline models’ performances (for example, the CLIP model in **Extended Figure 4b**), PLIP was able to effectively differentiate between normal colon mucosa (NORM) and colorectal adenocarcinoma epithelium (TUM), which both shared similar morphological and textural patterns.

**Figure 3.**
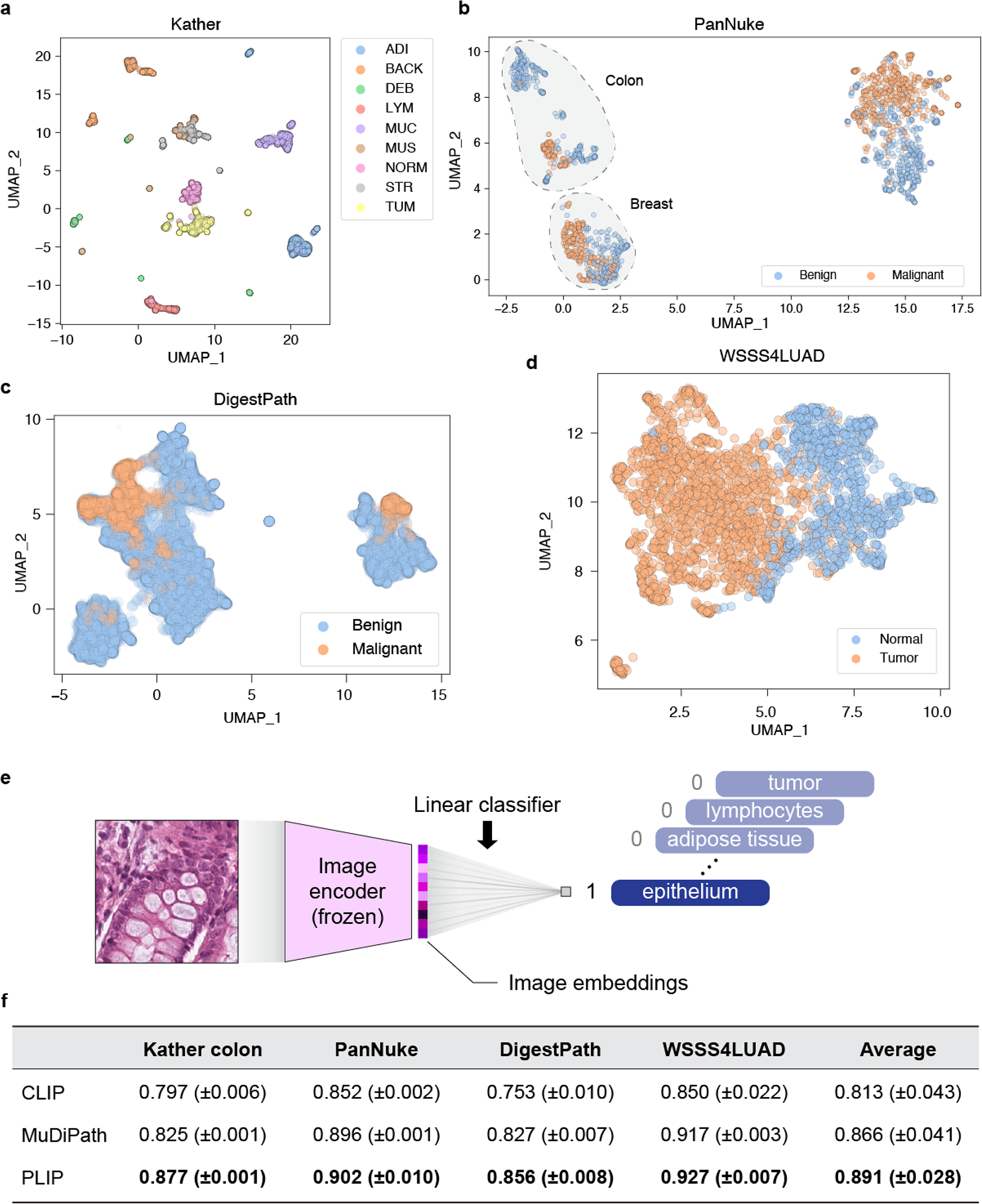
Image embedding analysis and linear probing results. **(a)** Image embeddings generated from the PLIP model in the Kather colon dataset. ADI: Adipose tissue, BACK: background, DEB: debris, LYM: lymphocytes, MUC: mucus, MUS: smooth muscle, NORM: normal colon mucosa, STR: cancer-associated stroma, TUM: colorectal adenocarcinoma epithelium. **(b)** Image embeddings generated from the PLIP model in the PanNuke dataset. **(c)** Image embeddings generated from the PLIP model in the DigestPath dataset. **(d)** Image embeddings generated from the PLIP model in the WSSS4LUAD dataset. **(e)** Graphical illustration of linear probing transfer learning. “Frozen” means the loss from the linear classifier will not be used to update the parameters of the image encoder. **(f)** F1 score in testing sets with mean (± standard deviation) from 5 repeated experiments with different random seeds. Best performed models are highlighted in boldface. The “Average” column shows the averaged performances across four datasets.

In the PanNuke dataset, PLIP revealed intriguing organ-specific separations, particularly highlighting breast and colon subsets. Indeed, the colon set formed two relatively clean sub-clusters, with one being enriched with the malignant tissue images (**Figure 3b**, **Extended Figure 5**). Furthermore, the PLIP model seemed able to separate normal/benign from tumoral/malignant image patches from the DigestPath and WSSS4LUAD datasets, respectively (**Figure 3c–d**, **Extended Figure 6**, **Extended Figure 7**). For DigestPath, we also noticed a clear separation between different image downsampling rates and staining variations (**Extended Figure 8**).

Encouraged by these findings, we hypothesize that the PLIP image encoder may serve as a preferred pre-trained backbone for various pathology image classification tasks. To this end, we trained a simple linear classifier on top of the image embedding vectors (**Figure 3e**) from the training splits of four different datasets (Kather colon, PanNuke, WSSS4LUAD, and DigestPath, see **Figure 2b**, **Extended Figure 2a**), and compared the classification performances on the testing splits of four datasets with two baseline models: image encoder from the original CLIP model and the multi-task pre-training of deep neural networks [34] (or “MuDiPath” for short), which was trained on a collection of 22 classification tasks with approximately 900k images that are not publicly available. For all the models, we extracted image embeddings from the last layer and used a linear classifier on top of these features. In the following, we formally abbreviate this adaptation to a new set of data by training the embedding/last hidden layer as “linear probing” [35].

By taking text description in natural language into account, our PLIP model sets itself apart from what others have done (*e.g.*, MuDiPath), and demonstrated superior performances across all four testing datasets (average macro F1: PLIP = 0.891, CLIP = 0.813, MuDiPath = 0.866, **Figure 3f**). In the Kather colon dataset with 9-class classification, PLIP achieved F1 = 0.877, surpassing the second-best model MuDiPath (F1 = 0.825) with a 6.30% improvement. In the PanNuke dataset with binary classification, PLIP achieved F1 = 0.902, which was the highest score among the compared models. In the DigestPath dataset, PLIP also improved by 3.5% in terms of F1 = 0.856 when compared to the second best model. Finally, PLIP achieved the highest F1 = 0.927 in the WSSS4LUAD datasets. Furthermore, an ablation study was conducted with different combinations of datasets, and the PLIP model was found to perform best when all OpenPath data (*Tweets+Replies+PathLAION*) were combined in training (**Extended Table 4**).

The results from the linear probing analysis suggest that PLIP, powered by the combined inferences of both images and text descriptions (especially in natural language), can achieve higher performances compared to traditional deep learning models trained solely with categorical labels. This is likely due to the rich content of the text annotations, which may enable the model to exploit higher-level semantic visio–linguistic relationships and provide a more comprehensive understanding of the images, including both visual and subvisual inter-cellular patterns.

### PLIP improves the retrieval of relevant pathology images from text input

By simultaneously learning both image and text representations, PLIP can identify and retrieve the most relevant images based on a given text or image input. To this end, a systematic analysis was conducted to evaluate the ability of text-to-image retrieval [36].

For text-to-image retrieval, we generated a vector representation for a searching query (in natural language) and identified the image that best matched the query (**Figure 4a**). This image search engine function is extremely useful in practice and may serve as a powerful educational resource. In this study, we collected four sets of images with captions of different lengths in terms of the number of words to evaluate the ability of pathology text-to-image retrieval: (i) Twitter validation dataset (Twitter, median caption length = 22); (ii) PathPedia images (PathPedia, median caption length = 12); (iii) PubMed pathology images [37] (PubMed, median caption length = 9); and (iv) pathology book collections [37] (Books, median caption length = 5) (**Figure 4b**). The Twitter validation dataset, which spanned 24 out of 32 hashtags, contained 2,023 paired image–text from 2022-11-16 to 2023-01-15 (**Figure 4c**, **Extended Figure 1b**), and was expected to have a similar image–text distribution to what the PLIP model had trained on. In contrast, PathPedia (Number of candidates for image retrieval = 210), PubMed (1,419 image–text pairs), and Books (558 image–text pairs) comprised relatively concise texts (**Figure 4d**).

**Figure 4.**
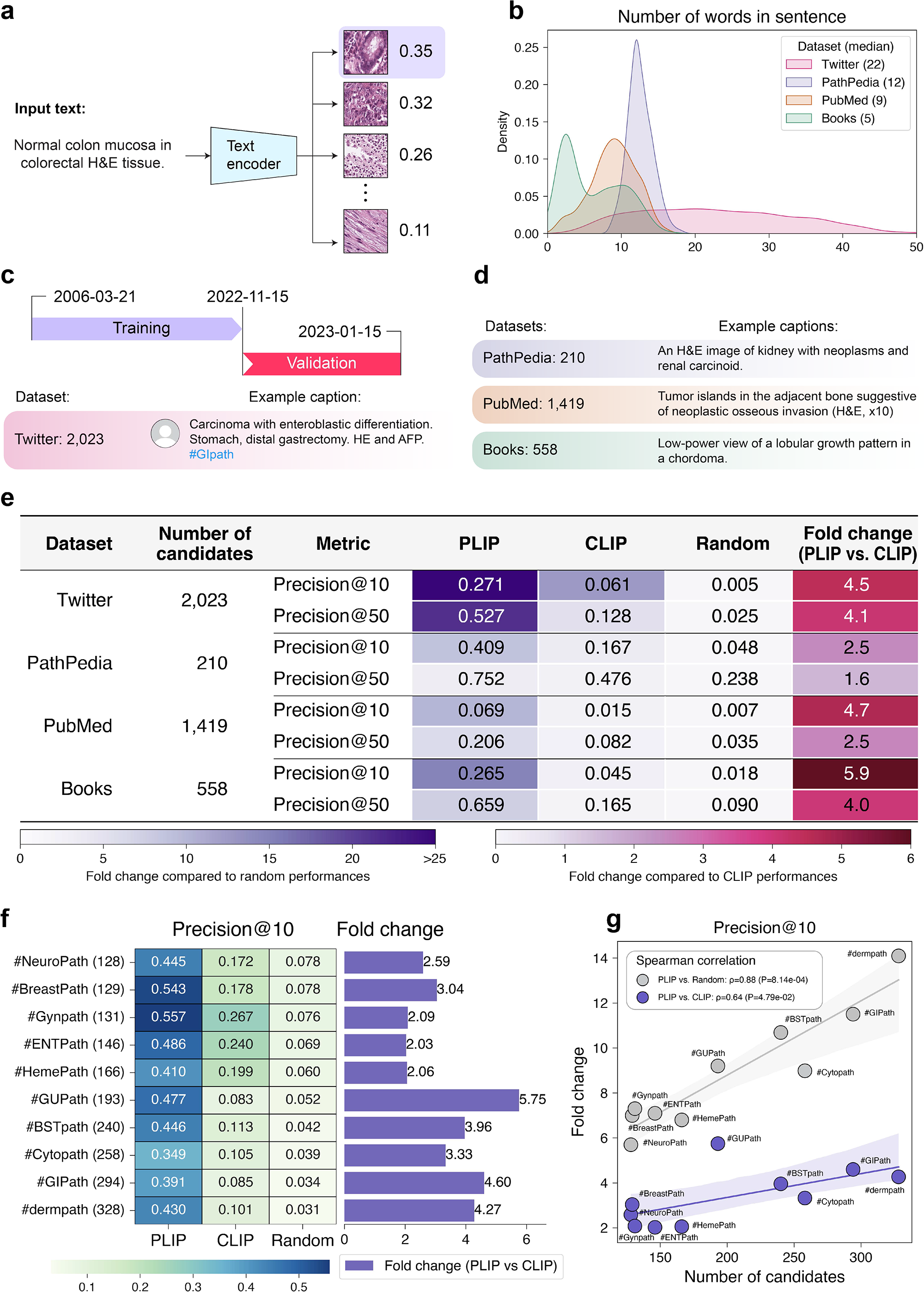
Text-to-image retrieval for pathology images. **(a)** Graphical illustration of pathology image retrieval from text input. **(b)** Density plot of the number of words per sentence across four validation datasets. **(c)** Description of the Twitter validation dataset and an example text caption. **(d)** Descriptions of the PathPedia, PubMed, and Books datasets and example text captions. **(e)** Image retrieval performances across validation datasets. **(f)** Text-to-image retrieval performances for Precision@10 within each of the pathology subspecialty-specific hashtags. **(g)** Spearman correlations between the number of candidates and fold changes for Precision@10 when comparing the PLIP model with CLIP and random, respectively.

We first employed the unseen portion of the Twitter data from 2022-11-16 to 2023-01-15 to evaluate the performance of image retrieval with two metrics: Precision@10 and Precision@50 – which refer to the precision of the target image being among the top 10 and top 50 retrieved images, respectively (See **Online Methods**). This task is very challenging; there are many images that could seem compatible with a given description (*e.g.*, “normal colon mucosa”) and thus finding the exact image associated with a given text can be difficult if there are other similar images. Nonetheless, we found PLIP improved the image retrieval significantly with Precision@10 = 0.271 (55.3x and 4.5x higher than random and CLIP results), and Precision@50 = 0.527 (21.4x and 4.1x higher than random and CLIP results) (**Figure 4e**). Given these sophisticated circumstances under a large pool of candidates (N = 2,023) and a diverse collection of various tissue types, the 52.7% chance to retrieve the target image within the top 50 makes the pathology image retrieval from only texts a challenging yet achievable task.

In addition, PLIP demonstrated a significant improvement in performance compared to both the baseline CLIP and random performances across different datasets (**Figure 4e**). In the PathPedia collection, PLIP achieved Precision@10 = 0.409 (8.6x and 2.5x higher than random and CLIP results), and Precision@50 = 0.752 (3.2x and 1.6x higher than random and CLIP results). In the PubMed pathology images collection, PLIP achieved Precision@10 = 0.069 (9.9x and 4.7x higher than random and CLIP results), and Precision@50 = 0.206 (5.9x and 2.5x higher than random and CLIP results). In the Books pathology images collection, PLIP achieved Precision@10 = 0.265 (14.8x and 5.9x higher than random and CLIP results), and Precision@50 = 0.659 (7.4x and 4.0x higher than random and CLIP results). Despite the varying numbers of candidates within each of the four image retrieval tasks, the fold change in terms of precision suggested that the Twitter validation dataset may benefit most from the PLIP model (fold change = 55.3 and 21.4 for Precision@10 and Precision@50 when comparing PLIP vs. Random). In contrast, the PathPedia image collection may have the least benefit from the PLIP model. These divergent performances observed across datasets are likely because the curated PathPedia collection does not cover all of the nuances and variations in the text for pathology images, leading to a difference in how humans describe them.

Motivated by the encouraging results of text-to-image retrieval on the Twitter validation dataset, we further conducted the text-to-image retrieval tasks for the Twitter validation dataset within each of the 24 available hashtags and presented the top 10 hashtags that had more than 100 available candidates (**Figure 4f–g**, **Extended Figure 9a–b**). This simulates the scenario when we want to retrieve the relevant pathology images given a known pathology subspecialty. When measured with Precision@10, we found that the gynecologic pathology (#Gynpath) may benefit most from the PLIP model, which had Precision@10 = 0.557 with 131 candidates and was 2.1x better than the CLIP performance (**Figure 4f**). When measured with Precision@50, head and neck pathology (#ENTPath) benefited most from the PLIP model, with Precision@50 = 0.925 with 146 candidates and was 1.6x better than the CLIP performance (**Extended Figure 9a**). Furthermore, the Spearman correlation analysis suggested that the performance improvements of the PLIP model were significantly correlated with the number of candidate images. For Precision@10, ρ = 0.88 (P-value = 8.14e-4) for PLIP vs. Random, and ρ = 0.64 (P-value = 4.79e-2) for PLIP vs. CLIP (**Figure 4g**). For Precision@50, ρ = 0.98 (P-value = 1.47e-6) for PLIP vs. Random, and ρ = 0.85 (P-value = 1.64e-3) for PLIP vs. CLIP (**Extended Figure 9b**). This observation suggested that the PLIP model exhibited a greater margin of improvement over baseline models when larger image pools were used for retrieval, indicating its potential for effective and robust retrieval under more complex scenarios. In summary, these results suggest that the fine-tuned PLIP model can effectively retrieve relevant pathology images with only text input.

### PLIP improves the retrieval of relevant pathology images from input image

Meanwhile, the ability of image-to-image retrieval [4] was evaluated by calculating the similarity between the image embedding of the target image and the image embeddings of the candidate images (**Figure 5a**). Evaluations were initially carried out from the Twitter validation dataset. As tweets often contain multiple images, we investigated whether using one image from a tweet can retrieve the rest of the other images from the same Twitter post (**Figure 5b**). Out of 2,023 image–text pairs from 925 tweets in the Twitter validation dataset, 525 tweets had at least 2 images, contributing to a total of 1,623 images to be searched. We assessed the retrieval ability by comparing each target image with all other images via the Precision@K, which measures the number of images in the top K retrieved that originate from the same Twitter post. The benchmark comparison was conducted by comparing PLIP with two baseline models: CLIP and MuDiPath. In addition, we included the SISH model [5], which had previously demonstrated superior performance in searching for whole slide images. Among the four models being compared, cosine similarity was used for measuring the similarity of the image embeddings generated from PLIP and CLIP, Euclidean distance was used for measuring the similarity of the image embeddings generated from MuDiPath, and Hamming distance was used for measuring the similarity of the image embeddings generated from SISH. The results presented in **Figure 5c** suggested that all four models were capable of retrieving relevant images, while the PLIP model achieved the best performance with Precision@10 = 0.646 (compared to CLIP at 0.353, MuDiPath at 0.336, and SISH at 0.356). Similarly, PLIP achieved the best performance with Precision@50 = 0.814 (compared to CLIP at 0.513, MuDiPath at 0.485, and SISH at 0.474). From the results, we found PLIP can achieve a relatively high Precision@10 and Precision@50 given more than 2,000 images in the pool for comparison whereas other models did not perform that well for Twitter image retrieval. The reason for this could be that although the images from the same Twitter posts may convey similar semantic knowledge, clinicians often post relatively different images in order to provide diverse perspectives (such as different magnification levels, different staining, etc.). This visual dissimilarity may have caused confusion in other models.

Additional evaluations were conducted on three external validation datasets, each with a distinct study focus (**Extended Figure 10**): (i) Tissue types: the Kather colon dataset (9 colon tissue types); (ii) Organ types: the PanNuke dataset (19 organs); and (iii) Staining textures: the KIMIA Path24C dataset (24 staining textures). By evaluating the class retrieval accuracy at top K, we can determine the purity of the retrieved K images from the same given class. It is worth noting that the score may decrease as K increases, as more images may be irrelevant in the top retrieval results. From the results, we found PLIP consistently outperformed other models across all three datasets (**Figure 5d–f**, **Extended Table 5**). For example, in the Kather colon dataset, we found PLIP achieved 0.998 when K = 10 (meanwhile, CLIP = 0.984, MuDiPath = 0.994, SISH = 0.993). In the PanNuke dataset, we found PLIP achieved 0.954 when K = 10 (meanwhile, CLIP = 0.915, MuDiPath = 0.927, SISH = 0.944). In the KIMIA Path24C dataset, we found PLIP achieved 0.906 when K = 10 (meanwhile, CLIP = 0.858, MuDiPath = 0.879, SISH = 0.885). These results under various testing scenarios including tissue types, organ types, and staining textures, suggested that PLIP is a preferred model to be used as an image-to-image retrieval system in pathology.

**Figure 5.**
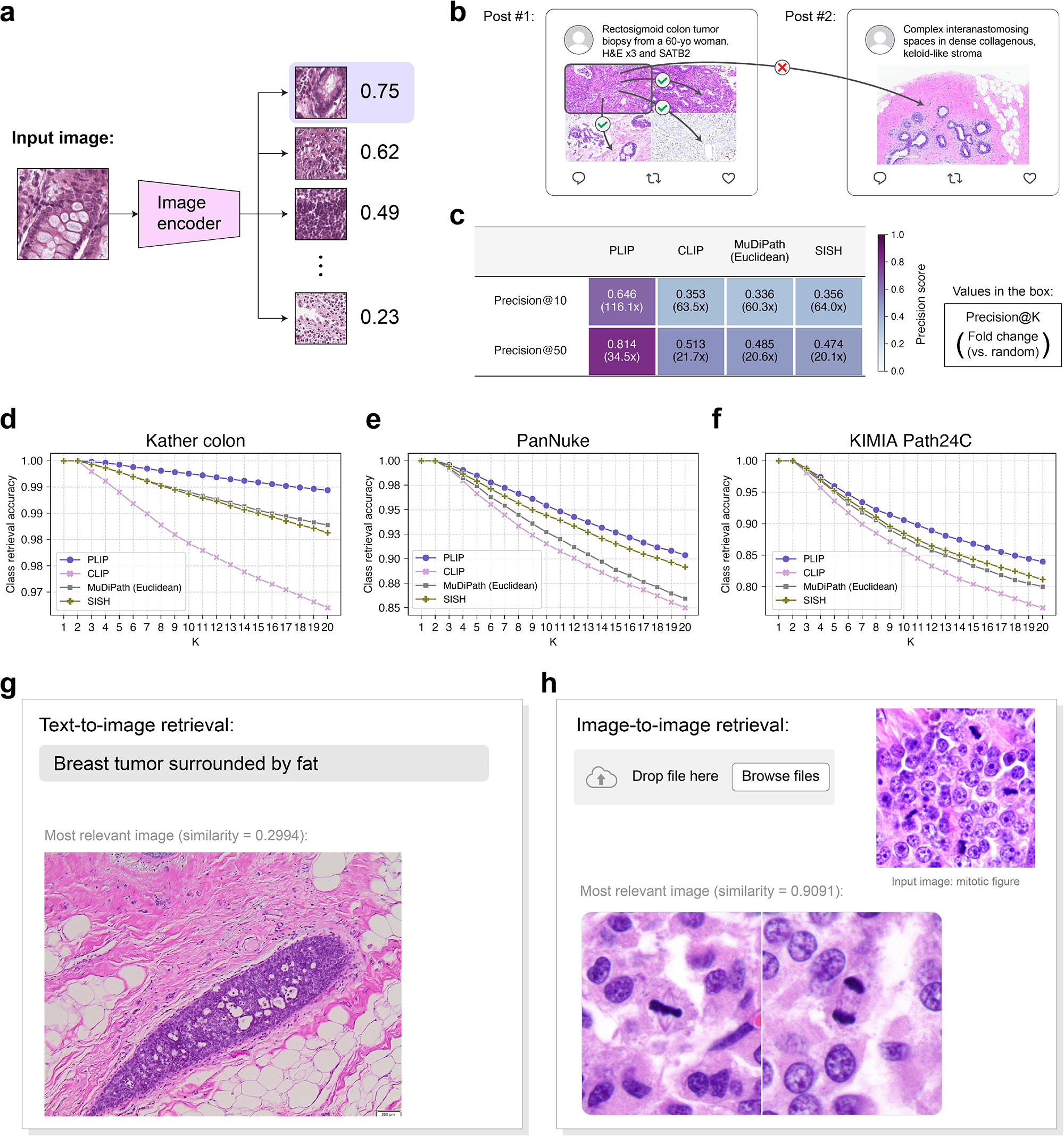
Image-to-image retrieval for pathology images. **(a)** Graphical illustration of image-to-image retrieval. **(b)** Illustration of image-to-image retrieval analysis on the Twitter validation dataset. **(c)** Image-to-image retrieval performances on the Twitter validation dataset. Values in boxes represent Precision@K score and the fold change compared to the random performance. **(d)** Image-to-image retrieval performances on the Kather colon dataset. **(e)** Image-to-image retrieval performances on the PanNuke dataset. **(f)** Image-to-image retrieval performances on the KIMIA Path24C dataset. **(g)** Examples of text-to-image retrieval. **(h)** Examples of image-to-image retrieval (featuring the mitotic figure).

Finally, the text-to-image and image-to-image retrieval systems can function as image search engines, enabling users to match images from multiple queries and retrieve the most relevant image based on a sentence description or an input image. As demonstrated and presented on our website (https://tinyurl.com/webplip), this generic system can comprehend semantic and interrelated knowledge, such as “Breast tumor surrounded by fat” (**Figure 5g**). This capability provides a powerful tool for exploring and retrieving large pathology datasets, allowing users to efficiently and accurately identify relevant images that meet specific criteria. Additionally, the ability of image-to-image retrieval was further demonstrated and can be used to retrieve relevant pathology images similar to the target image input, for example, images that contain mitotic figures, demonstrating its ability to comprehend the key concepts from the input image (**Figure 5h**).

## Discussion

With the unprecedented evolution of machine learning in both computer vision and natural language processing (NLP), the cornerstone of establishing a model for general purposes has been heavily reliant on high-quality annotated data. Unlike other fields, annotating pathology images is extremely expensive and laborious, which requires a high level of domain expertise and years of professional education [38,39]. As a consequence, the shortage of annotated pathology images has limited the progress of artificial intelligence in digital pathology, making it a relatively understudied area [40]. Unfortunately, this obstructs the advancement of AI to understand and interpret histopathological features, decipher anatomy and disease heterogeneity, and eventually contribute to precision medicine [40].

The explosion of data on the Internet has drawn significant attention yet is underutilized by medical AI communities. Especially on social media, the activity of crowdsourcing, spontaneous, and interactive knowledge sharing has opened up many opportunities for data-hungry fields [32,41–43]. As one of the largest public social networks, Twitter has become the most proactive community for pathologists [16]. Despite the existence of vast amounts of unorganized data that give rise to difficulties in obtaining images, Twitter hashtags render a relatively organized and finite subset within a particular topic [12], which can be a valuable resource for AI to study pathology on a large scale. This paired image–text collection provides unique structural knowledge and may help AI to comprehend both global and local pathological features. In this study, leveraged by our OpenPath dataset that is now publicly available, we developed our PLIP model by fine-tuning the state-of-the-art model in visual–language representation and learning [27].

As a novel AI model, PLIP can enable significant strides in the field of pathology by employing joint vision and language learning and inference. Through four different learning tasks including zero-shot learning, linear probing, text-to-image retrieval, and image-to-image retrieval, PLIP demonstrated its superior performance compared to previous state-of-the-art baselines for each of these tasks.

One of our key algorithmic contributions was the development of an improved image representation for pathology images, which facilitates numerous downstream tasks such as zero-shot learning and linear probing. Unlike other canonical machine learning approaches in digital pathology that learn from a fixed set of labels, our PLIP model is a general solution that can be applied to a wide range of tasks, including adaptation to a new set of data and provide zero-shot predictions given any text inputs. This capability of handling a varying number of classes is particularly valuable in scenarios where the learning objectives change after the model has been trained. Furthermore, this zero-shot ability also aligns with the constantly evolving criteria for diagnosis in pathology [8]. The improved ability of image representation was then quantitatively demonstrated by linear probing analysis. Our current linear probing evaluation benchmarks, while not directly comparable due to the fact that MuDiPath has been trained on a private dataset [34], provide so far the closest match to other studies in this field.

The results of the linear probing ablation study indicate that incorporating the most-liked replies can lead to a significant improvement in performance. This finding highlights the potential use of conversations among pathologists as a valuable source of knowledge. In addition to the most-liked replies, the discrepancies among all replies/answers in a Twitter post can also indicate the level of consensus or disagreement among pathologists, which may provide insights into the level of difficulty involved in diagnosing a pathology image, making it a valuable area for future research.

Unlike searching keywords and sentences from Google and indirectly matching the images from the target text, our proposed pathology image retrieval allows direct comparison between input sentences and images. This innovation in the field of pathology has the potential to serve as a powerful educational resource for trainees and could also be used as an information retrieval system in clinical workflows. While previous work has made achievements on pathology image-to-image retrieval [4,5,44–48], there is a limited literature on text-to-image retrieval in pathology due to the lack of paired image–text data. For example, in [49], the authors implemented a text-to-image retrieval model for bladder cancer samples, which was trained on only 221 whole slide images. In our study, the proposed foundation AI model can perform text-to-image searching in pathology across various subspecialties, which greatly enhances the interaction between humans and AI.

This study has several limitations. First, despite rigorous cohort exclusion criteria and a stringent quality control pipeline applied to the proposed dataset, the quality of the image–text pairs may still contain some irrelevant data. Second, although several image pre-processing and transformation algorithms were applied to the image encoder upfront, identifying and accounting for the various magnifications of pathology image patches and different staining styles still remained a challenging and unsolved problem. However, encouraged by our current results, it is reasonable to conclude that the PLIP model already has the potential to recognize and adapt to images at various magnifications and staining protocols. Third, it is worth noting that zero-shot classification using prompts can be difficult to stabilize, as even a slight variation in the prompt could alter the results by a large margin [50]. We expect that continued endeavors to optimize prompts may lead to further improvements in zero-shot performance. Moreover, while the text-to-image retrieval results presented promising performances and large improvements over baseline models, the retrieval datasets used were limited in size owing to the limited literature available on this topic, and current results do not suggest that search can be replaced entirely by PLIP. As a challenging yet meaningful technique, the image retrieval task requires larger datasets of images and captions to better understand the effectiveness of the model. Lastly, limited by computational ability, all input images were resized to 224x224 pixels according to the original CLIP, which might lose some visual and subvisual patterns in pathology images.

Our PLIP model and the achievements on various learning tasks were made possible due to the establishment of the largest publicly available dataset, OpenPath, containing paired pathology images and text descriptions. With these findings and the availability of our dataset and model, we anticipate that the field of AI in pathology can benefit from OpenPath and PLIP, and facilitate data-centric AI in pathology.

## Online Methods

### Description of the OpenPath dataset

#### Release Policy

In accordance with the policy and regulation of Twitter and other entities including LAION, all information provided in the datasets is linked to the original source of the data. Specifically, data that has been collected from Twitter is released in the form of Tweet IDs, and data that has been collected from LAION is released in the form of URLs to images. Interested users will need to refer to the original sources to understand if their usage is compliant with the policies and regulations.

#### Twitter collection

All Twitter posts (tweets) with English captions under 32 pathology subspecialty-specific hashtags were included according to the recommendation from the 2016 United States and Canadian Academy for Pathology (USCAP) meeting and the Pathology Hashtag Ontology projects (https://www.symplur.com/healthcare-hashtags/ontology/pathology/). The complete list of hashtags and descriptions was presented in **Extended Table 1**. Tweets were collected from 2006-03-21 (the date of the first Twitter post) to 2022-11-15. Conversations including replies were collected for each of the tweets from 2006-03-21 to 2022-11-22 (7 days after the last tweet was collected on 2022-11-15). In total, we collected 232,067 tweets and 243,375 image–text pairs. Among those tweets, we further collected 88,250 replies which (1) the associated tweets had replies, (2) sentences contained at least one keyword from the ICD11 codebook (version 2022-02), and (3) received the highest number of likes among all replies.

Several cohort exclusion criteria were applied to our raw dataset (**Extended Figure 1a**), which include (1) tweets that were marked as possibly sensitive by Twitter; (2) duplicate image–text pairs; (3) images that cannot be downloaded or opened; (4) non-pathology images by pathology image classification model; and (5) texts which had question marks since questions often do not contain information about the image. After our stringent data inclusion and exclusion criteria, we ended up with 116,504 unique image–text pairs from tweets, and 59,869 unique image–text pairs from the top replies. As part of the image retrieval tasks, the Twitter validation dataset was further collected from 2022-11-16 to 2023-01-15 with the exact same cohort inclusion and exclusion criteria (**Extended Figure 1b**), and resulted in 2,023 unique image–text pairs.

#### PathLAION collection

We established the PathLAION collection, which contains the pathology image–text data from sources beyond Twitter on the Internet. PathLAION is a subset of pathology images from the LAION-5B dataset, which originally contained 5.85 billion image–text pairs from the Internet [18]. This PathLAION collection was obtained by feeding a pathology image from the Twitter dataset to LAION-5B to retrieve the 5,000 most similar images, measured by the cosine similarity of CLIP image embeddings. The subsampling process stopped once all images retrieved were duplicated. By repeating this step with 1,000 images sampled from the Twitter dataset, we obtained 32,041 unique pathology images that were not present in the Twitter dataset.

#### Quality control of the training dataset

An additional quality control pipeline was applied to the established OpenPath dataset. A pathology image classifier was developed to exclude non-pathology images from both the Twitter and LAION datasets. As not all downloaded images were microscopic pathology images, this step was necessary to ensure the quality and relevance of the dataset.

To ensure the high-quality text descriptions of the associated images, the texts of the tweets and replies were cleaned using the following pipeline: (1) remove “@username” in the sentence; (2) remove hashtags “#”; (3) remove HTML tags for italic and bold formatting (for example, replace “ *keyword*” or “keyword” with “keyword”). (4) remove emojis; (5) remove the symbols of newline (\n) and carriage return (\r). (6) remove extra white spaces; (7) remove links start with “http://” and “https://”. Furthermore, tweets containing question marks were removed, as they are often used by practitioners to inquire about a pathology image rather than provide informative descriptions. For PathLAION, all image–text pairs with non-English captions were removed by using LangDetect (https://pypi.org/project/langdetect/). The complete statistics of text lengths in the final training dataset are presented in **Extended Table 3**.

### External validation datasets

To evaluate the performance of the proposed machine learning algorithm, a set of publicly available datasets were gathered.

For zero-shot and linear probing analysis, four datasets were collected: (1) the Kather colon dataset [24], which consisted of 100,000 training image patches and 7,180 validation image patches from the TCGA colorectal H&E images. (2) the PanNuke dataset [28], which contained 19 organ types with 5 different types of nuclei (neoplastic, inflammatory, connective tissue, epithelial, and dead), had a total of 7,558 image patches that contained at least 1 cell. Among them, images were deemed to be “malignant” if both the total number of neoplastic cells >= 10 and occupied over 30% of the total cells. Alternatively, images were deemed to be “benign” if no neoplastic cells exist. This resulted in 2,866 malignant images and 3,368 benign images. (3) DigestPath [29], initially from low-magnification whole slide images, were then cropped into multiple patches with various downsampling rates (2x, 4x, 8x, 16x, and 32x) with pixel sizes = 224x224 and 10% patch overlapping. Background patches were excluded if more than 50% of the pixels contained no tissue (RGB threshold = 200). With pixel-level annotation of malignancy, images were deemed to be malignant if the region of malignancy occupied more than 50% of the tissue area within an image patch. As a result, 6,690 malignant images and 56,023 benign images were obtained. (4) For WSSS4LUAD [30], all images were binarized to either tumor or normal, based on the existence of tumors on an image. This resulted in 6,579 tumor images and 3,512 normal images. All image patches were then resized to 224x224 before feeding into the image encoders. Except for the Kather colon dataset which had pre-existing training and validation splits, all other datasets (PanNuke, DigestPath, and WSSS4LUAD) were randomly divided into training and validation sets with a ratio of 70%:30%. To prevent data leakage on DigestPath, training and validation splits were separated according to their unique sample ID. To ensure the benchmarks are comparable, both zero-shot and linear probing analyses were evaluated on the validation split.

For text-to-image retrieval analysis, we collected three datasets in addition to the Twitter validation dataset. These datasets include PathPedia (http://www.pathpedia.com/) as well as the pathology collections of PubMed [37] and Books [37]. For PathPedia, we subset the original set of data by dropping duplicated captions, which resulted in 210 unique image–text pairs. In PathPedia, captions were further curated with “An image of Category with Subclass”, where the Category refers to the organ type (*e.g.*, kidney) and the Subclass refers to a specific disease subtype (*e.g.*, neoplasm). For PubMed and Books, duplicated captions were removed. Captions containing more than 100 characters or less than 3 characters were considered outliers and excluded from our subsequent analysis. In addition to this, looking at the captions we realized that often most of the information is contained in the first sentence. To this end, we removed all the text after the first period and finally checked again if the caption is between 3 and 100 characters.

For image-to-image retrieval analysis, we used the Kather colon dataset with 9 different tissue types and the PanNuke dataset with 19 different organ types. In addition, we included the KIMIA Path24C dataset [51] with 24 different pathology image staining textures to evaluate the model’s ability to retrieve images with the same textural pattern.

### Model training and tuning

All experiments were run in python 3.9. Detailed software versions are: pytorch v1.13, CUDA v11.7, CUDNN v8.5.0, scipy v1.9.3, torchvision v0.14.0, pillow v9.1.0, scikit-learn v1.1.2, scikit-image v0.19.2, pandas v1.4.2, numpy v1.23.5, and multiprocess v0.70.13.

#### Model training

We established our PLIP model with the same architecture described in [27]. This architecture is based on a vision transformer as the image encoder (ViT-32-Base, which can take in input images of size = 224 by 224 pixels), and a text transformer as the text encoder (with max sequence length equal to 76 tokens) [52]. Images were initially resized to a maximum dimension of 512 pixels, then were randomly cropped to 224 by 224 pixels in size before feeding into the image encoder. Both image and text encoders output vectors of 512 dimensions, and were optimized by minimizing the contrastive loss on a given batch [53]. Contrastive learning imposes a higher cosine similarity in paired image and text, forcing the model to learn the correct relationships between images and texts.

To find the optimal set of hyper-parameters, different combinations of training datasets and learning rates were searched on the linear probing task. With the model being evaluated on every quarter of an epoch (1 step = 1/4 epochs) given a total of 12 epochs, we found the best-performed model was trained with optimal learning rate = 1e-5, steps = 10, and using all training datasets (*Tweets+Replies+PathLAION*). An ablation study was conducted with different combinations of datasets and the same hyper-parameters, and the results shown in **Extended Table 4** indicate that the combination of all data (*Tweets*+*Replies*+*PathLAION*) allowed us to get the best-performed PLIP model.

#### Zero-shot classification

Since images and texts are encoded into the same vector space, the proposed model has the ability to learn the similarity between a target image with text candidates, and the text prediction is the one with the highest similarity. Also known as zero-shot learning [19], this ability to transfer a new unseen dataset of images and texts requires zero training data. In this study, we adapted four datasets (Kather colon, PanNuke, DigestPath, and WSSS4LUAD) with their validation splits to evaluate the ability of zero-shot classification performances. The text candidates were generated according to their tissue type. In the Kather colon dataset, we generate the sentences with “An H&E image of {keyword}.”, where the keyword can be (1) adipose tissue, (2) background, (3) debris, (4) lymphocytes, (5) mucus, (6) smooth muscle, (7) normal colon mucosa, (8) cancer-associated stroma, and (9) colorectal adenocarcinoma epithelium. In the PanNuke and DigestPath datasets, we generate the sentences with “An H&E image of {keyword} tissue.”, where the keywords can be (1) benign and (2) malignant. In the WSSS4LUAD dataset, we generate the sentences with “An H&E image of {keyword} tissue.”, where the keywords can be (1) normal and (2) tumor. To determine the confidence interval, zero-shot predictions were bootstrapped 100 times, each with 70% of the data used in the calculation.

#### Linear probing

Linear probing is a commonly used technique for assessing the quality of the features extracted by a model [35,54]. In this study, feature embeddings were extracted from the images and then a linear classifier was trained on top of these embeddings for a given classification task. To accomplish this, we used a logistic regression classifier from the SGDClassifier module in the sklearn python package [55]. For benchmark comparison, we compared our PLIP model backbone to the baseline CLIP model backbone, as well as the multi-task pre-training deep neural networks model [34] (or MuDiPath for short) with DenseNet121 architecture [56]. We evaluated the performance of the linear classifier using L2 regularization with various regularization multipliers (⍺ = 0.1, 0.01, 0.001, and 0.0001) on the validation splits for all models. The best-performing linear classifier was selected based on the average performance from the results generated by five different random seeds (seed = 1, 2, 3, 4, 5). The linear probing analysis was also used to determine the most suitable PLIP model for other experiments.

#### Image retrieval

Similar to zero-shot learning, which can identify the closest text from a pool of candidates given an image, image retrieval is a technique that can identify the closest image from a pool of candidates given a text [36] or image [4]. This was accomplished by directly calculating the cosine similarity of each paired image–text or image–image under the same embedding space.

The performance of text-to-image retrieval was evaluated by two metrics: Precision@10 and Precision@50 – which refers to the precision of the target image being among the top 10 and top 50 retrieved images, respectively [57]. For example, given a set of 500 image–text pairs and each of the text was fed into the image retrieval task to find the corresponding image. If this process is repeated for all 500 pairs and 60% (or 300 images) of the true positive images can be found under the top 10 closest matches from the pool of 500, then the precision@10 becomes 0.60.

The performance of image-to-image retrieval was evaluated by class retrieval accuracy across models, which is also known as the mean average precision at K (MAP@K) [5]. We consider the retrieved image “relevant” if the class of the retrieved image is the same as the target input, and the precision is the fraction of relevant images among the top K retrieved images. The average precision@K (AP@K) is defined as

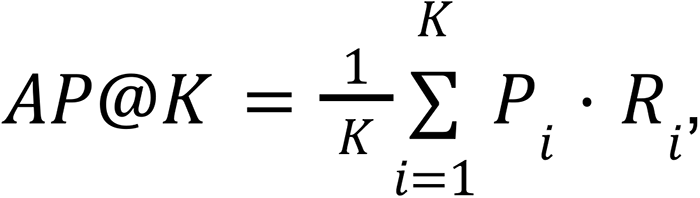

where *P_i_* is precision at *i*, and *R*_i_ is the relevant inidcator at *i*(*R*_i_ = 1 if the *i*th item is relevant, *R_i_* = 0 if not). The final class retrieval accuracy was calculated by

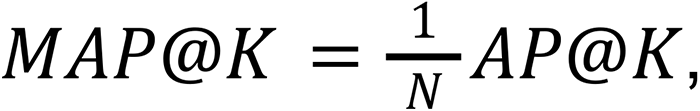

where *N* is the total number of samples. In our image-to-image benchmark comparison, cosine similarity was adopted to compare the similarity between image embeddings for PLIP and CLIP models, while MuDiPath and SISH model [5] calculated the similarity between image embeddings by Euclidean distance and Hamming distance, respectively. According to the guideline for the SISH model, we employed a large vector quantized VAE encoder based on their pre-trained weights.

### Evaluation metrics and statistical analysis

The F1 score was used to evaluate the performances of zero-shot and linear probing. Ranges from 0 – 1, the F1 score is calculated from the harmonic mean of precision and recall score:

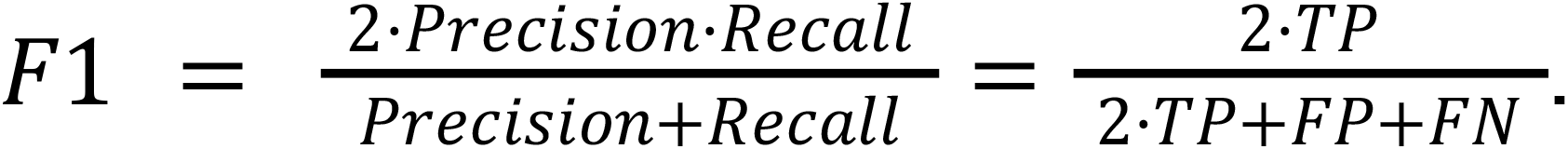

While the higher the better, it represents the overall performance of a model on a classification task. To evaluate the performance of the image retrieval task,

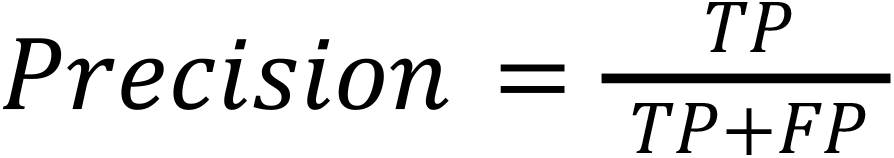

was adopted to quantitatively measure the fraction of the target image that exists among top *K* retrieved images. In our image retrieval experiments, we evaluated precision with *K* = 10 and *K* = 50.

Spearman correlation with two-sided P-value was used to evaluate the correlation between the number of candidates and fold changes for the image retrieval task.

## Data Availability

All data in OpenPath is publicly available from Twitter and LAION-5B (https://laion.ai/blog/laion-5b/). Twitter IDs used for training and validation were provided in the supplementary file. Validation datasets are all publicly available and can be accessed from: Kather colon dataset (https://zenodo.org/record/1214456), PanNuke (https://jgamper.github.io/PanNukeDataset/), DigestPath (https://digestpath2019.grand-challenge.org/), WSSS4LUAD (https://wsss4luad.grand-challenge.org/), PathPedia (http://www.pathpedia.com/), PubMed and Books pathology collection (https://warwick.ac.uk/fac/cross_fac/tia/data/arch), KIMIA Path24C (https://kimialab.uwaterloo.ca/kimia/index.php/pathology-images-kimia-path24/). The trained model and interactive results can also be visualized at https://tinyurl.com/webplip.

## Code Availability

The trained model and source codes can be accessed at https://tinyurl.com/webplip.

## Acknowledgements

F.B. is supported by the Hoffman–Yee Research Grants Program and the Stanford Institute for Human-Centered Artificial Intelligence. J.Z. is supported by the Chan-Zuckerberg Biohub.

