## Supplementary Information for "Leveraging medical Twitter to build a visual–language foundation model for pathology AI"

### Extended Tables and Figures

**Extended Table 1**. List of pathology hashtags on Twitter used in this study from 2006-03-21 (the date of the first Twitter post) to 2022-11-15 and the original number of data before data filtering.

| | Hashtag | Text | # tweets | # tweets with images | # images |
|---|---|---|---|---|---|
| 1 | Autopsy | Autopsy Pathology | 33804 | 7287 | 8444 |
| 2 | BloodBank | Blood Banking & Transfusion Medicine | 16030 | 6064 | 7996 |
| 3 | blooducation | Blood Banking & Transfusion Medicine | 10574 | 2968 | 3264 |
| 4 | BreastPath | Breast Pathology | 5999 | 3883 | 8436 |
| 5 | BSTpath | Bone And Soft Tissue Pathology | 8361 | 5680 | 13839 |
| 6 | CardiacPath | cardiovascular pathology | 1788 | 1104 | 1788 |
| 7 | ClinPath | Clinical Pathology | 1083 | 487 | 599 |
| 8 | Cytopath | Cytopathology | 13036 | 9005 | 16531 |
| 9 | dermpath | Dermatopathology | 30612 | 21307 | 42434 |
| 10 | EndoPath | Endocrine Pathology | 1889 | 1004 | 1997 |
| 11 | ENTPath | Head And Neck Pathology | 6342 | 4506 | 9541 |
| 12 | EyePath | Ophthalmic Pathology | 908 | 621 | 1135 |
| 13 | FNApath | Fine Needle Aspirate (FNA) Cytopathology | 361 | 295 | 565 |
| 14 | ForensicPath | Forensic Pathology & Forensics | 1042 | 464 | 719 |
| 15 | GIPath | Gastrointestinal and Liver Pathology | 17214 | 11985 | 27284 |
| 16 | GUPath | Genitourinary Pathology | 15216 | 9500 | 19819 |
| 17 | Gynpath | Gynecologic Pathology | 7543 | 5378 | 11597 |
| 18 | HemePath | Hematopathology | 18395 | 11088 | 20671 |
| 19 | HPBpath | Hepatobiliary pathology | 128 | 59 | 122 |
| 20 | IDpath | Infectious Disease Pathology | 733 | 556 | 1063 |
| 21 | MolDx | Molecular Pathology | 1502 | 788 | 959 |
| 22 | nephpath | Nephropathology | 207 | 130 | 240 |
| 23 | NeuroPath | Neuropathology | 9978 | 4424 | 9052 |
| 24 | OralPath | Oral Pathology | 2083 | 1403 | 2813 |
| 25 | pancpath | Pancreatic pathology | 362 | 187 | 429 |
| 26 | PathGME | Pathology Graduate Medical Education | 19 | 11 | 14 |
| 27 | pathInformatics | Pathology Informatics | 158 | 92 | 134 |
| 28 | patientbloodmanagement | Blood Banking & Transfusion Medicine | 1752 | 504 | 561 |
| 29 | PediPath | Pediatric Pathology | 8028 | 4424 | 9072 |
| 30 | PulmPath | Pulmonary And Pleural Pathology | 6611 | 3891 | 8430 |
| 31 | RenalPath | Renal and Medical Kidney Pathology | 5933 | 3738 | 6563 |
| 32 | SurgPath | Surgical Pathology | 4376 | 3405 | 7264 |
| | | **Total** | 232067 | 126238 | 243375 |

**Extended Table 2**. Quality assessment of OpenPath dataset by manual inspection. A total of 1,000 images were subsampled from 10 batches with seed = {0, 1, …, 9}, and each batch contained 100 images.

| Category | Image types | Number of images | Percentile (%) |
| --- | --- | --- | --- |
| Pathology-related images | High-quality whole slide image patches | 945 | 94.5% |
| | PPT/presentation with pathology images | 36 | 3.6% |
| | Webpage/screenshot with pathology images | 4 | 0.4% |
| Other non-pathology images | PPT/presentation/screenshot without pathology images | 5 | 0.5% |
| | Pathology illustration images | 3 | 0.3% |
| | Tissue mass | 3 | 0.3% |
| | Glass slide photos | 2 | 0.2% |
| | Other images (radiology images, etc.) | 2 | 0.2% |
| | **Total** | 1,000 | – |

**Extended Table 3**. Statistics of text length in the OpenPath training dataset.

| Training dataset | Number of words | | | Number of characters | | |
|---|---|---|---|---|---|---|
| | Median | Min | Max | Median | Min | Max |
| *Tweets* | 20 | 2 | 56 | 138 | 12 | 296 |
| *Replies* | 15 | 1 | 57 | 98 | 6 | 295 |
| *PathLAION* | 9 | 1 | 347 | 84 | 5 | 2,277 |
| *All combined* | 17 | 1 | 347 | 118 | 5 | 2,277 |

**Extended Table 4**. Ablation study on linear probing with different combinations of training datasets across four validation datasets. Performances are reported as mean F1 scores with standard deviation in parentheses. Best performing scores are highlighted in boldface.

| Training dataset | Linear probing validation datasets | | | | |
|---|---|---|---|---|---|
| | Kather colon | WSSS4LUAD | DigestPath | PanNuke | All |
| *PathLAION* | 0.857 (±0.005) | 0.886 (±0.007) | 0.797 (±0.025) | 0.866 (±0.009) | 0.851 (±0.036) |
| *Tweets* | 0.764 (±0.006) | 0.917 (±0.012) | 0.848 (±0.012) | **0.907 (±0.004)** | 0.859 (±0.061) |
| *Tweets+PathLAION* | 0.833 (±0.008) | 0.906 (±0.007) | 0.831 (±0.015) | 0.880 (±0.015) | 0.863 (±0.033) |
| *Tweets+Replies+PathLAION* | **0.877 (±0.002)** | **0.927 (±0.007)** | **0.856 (±0.009)** | 0.902 (±0.011) | **0.891 (±0.028)** |

**Extended Table 5**. Image-to-image retrieval performances for the Kather colon dataset (9 colon tissue types), the PanNuke dataset (19 pathology subspecialties), and the KIMIA Path24C dataset (24 staining textures). Performances were reported in by class retrieval accuracy when K = 5, 10, 15, and 20. Best performances were highlighted in boldface.

| Dataset (Number of classes) | K | Model | | | |
|---|---|---|---|---|---|
| | | PLIP | CLIP | MuDiPath | SISH |
| Kather colon (9) | 5 | **0.999** | 0.994 | 0.998 | 0.998 |
| | 10 | **0.998** | 0.984 | 0.994 | 0.993 |
| | 15 | **0.996** | 0.978 | 0.991 | 0.990 |
| | 20 | **0.994** | 0.972 | 0.988 | 0.986 |
| PanNuke (19) | 5 | **0.985** | 0.966 | 0.974 | 0.979 |
| | 10 | **0.954** | 0.915 | 0.927 | 0.944 |
| | 15 | **0.927** | 0.879 | 0.889 | 0.915 |
| | 20 | **0.904** | 0.850 | 0.859 | 0.892 |
| KIMIA Path24C (24) | 5 | **0.960** | 0.936 | 0.951 | 0.952 |
| | 10 | **0.906** | 0.858 | 0.879 | 0.885 |
| | 15 | **0.868** | 0.804 | 0.833 | 0.844 |
| | 20 | **0.840** | 0.766 | 0.800 | 0.812 |

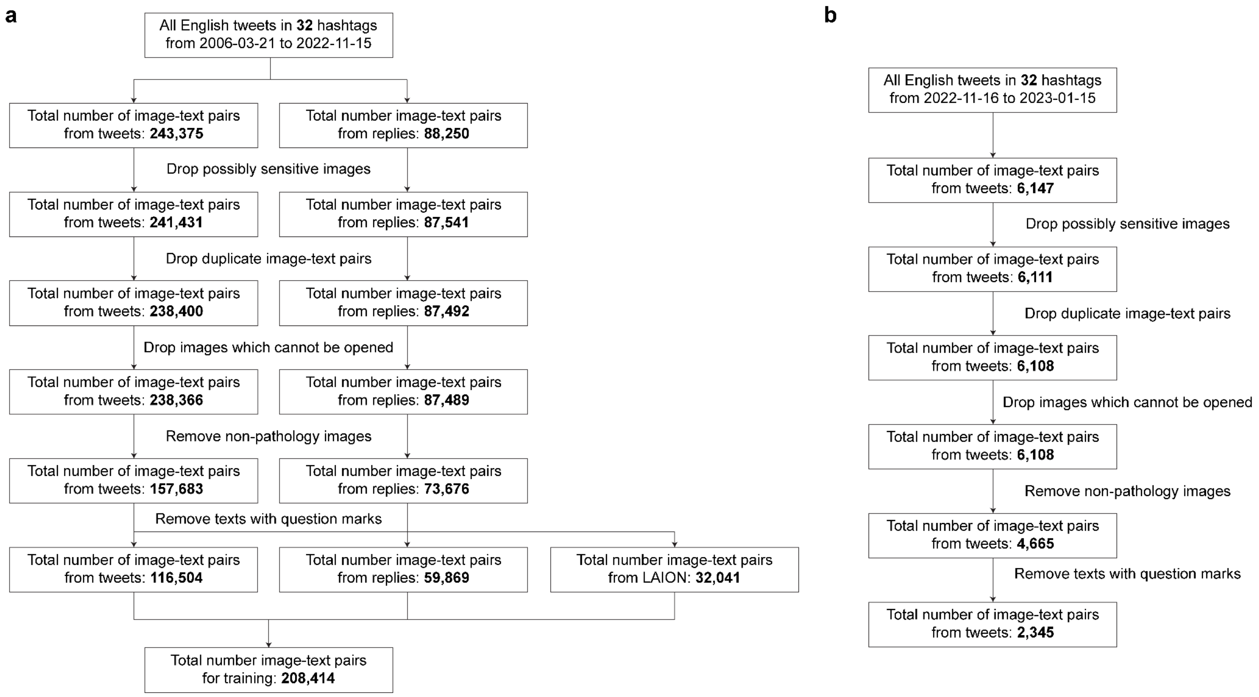


**Extended Figure 1**. Cohort inclusion and exclusion criteria and flowcharts. (**a**) Twitter training dataset from 2006-03-21 to 2022-11-15. (**b**) Twitter validation dataset for image retrieval from 2022-11-16 to 2023-01-15.

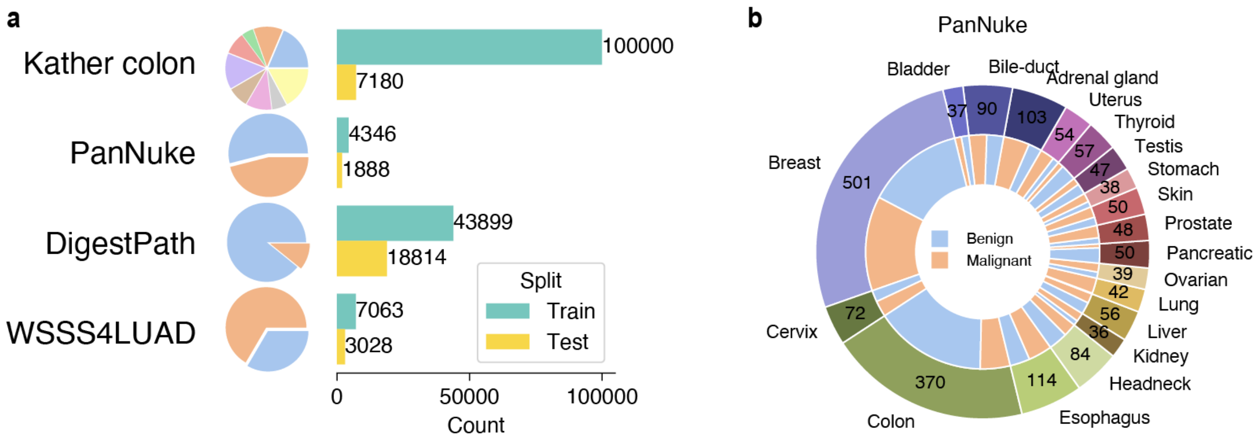


**Extended Figure 2**. Datasets for zero-shot classifcation and linear probing. (**a**) Training and testing splits. (**b**) PanNuke dataset with 19 different tissue types.

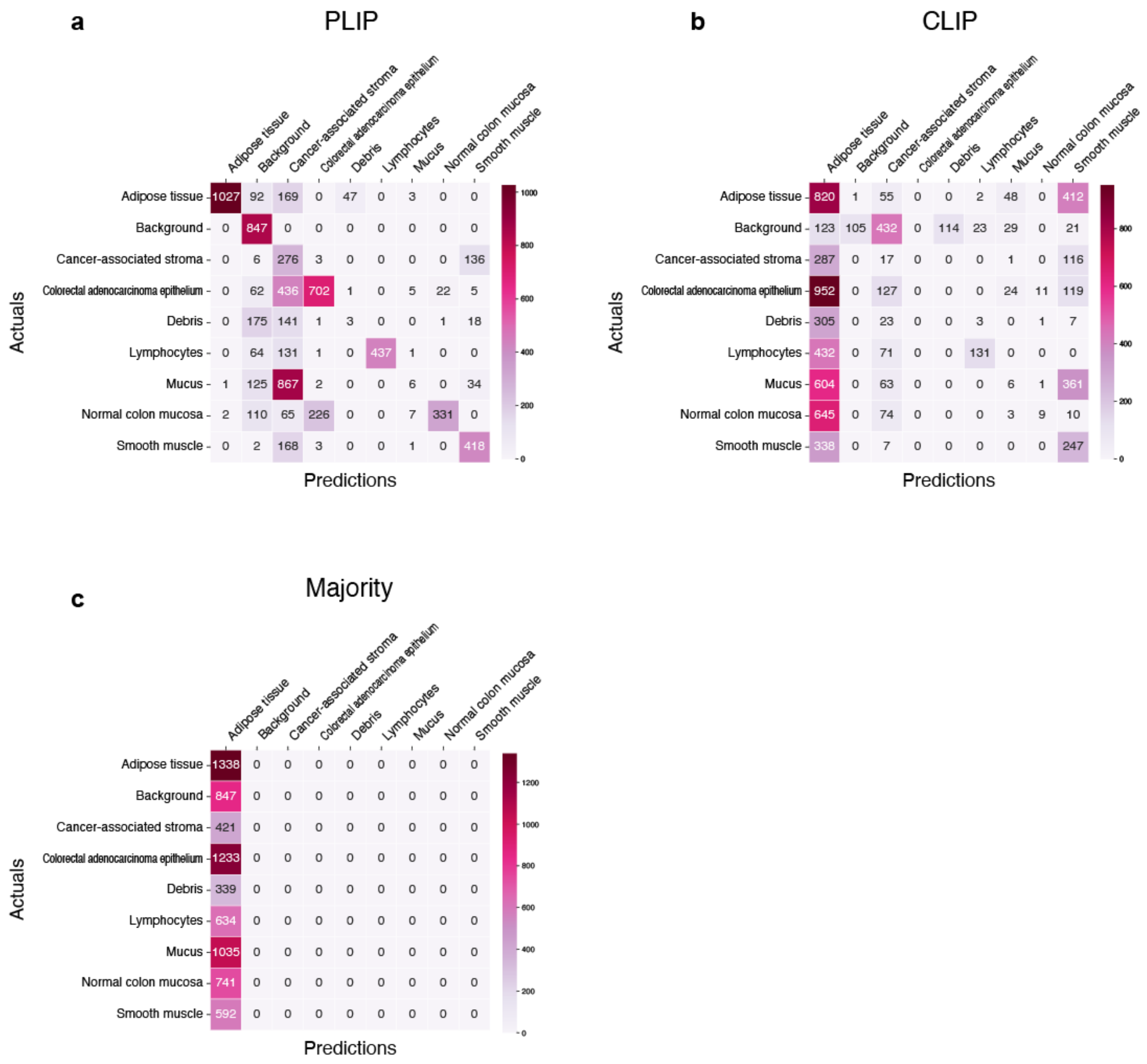


**Extended Figure 3**. Confusion matrix from zero-shot learning in the Kather colon dataset. (**a**) Confusion matrix of PLIP model; (**b**) Confusion matrix of CLIP model; (**c**) Confusion matrix of the results from predicting the majority class (or Majority in short).

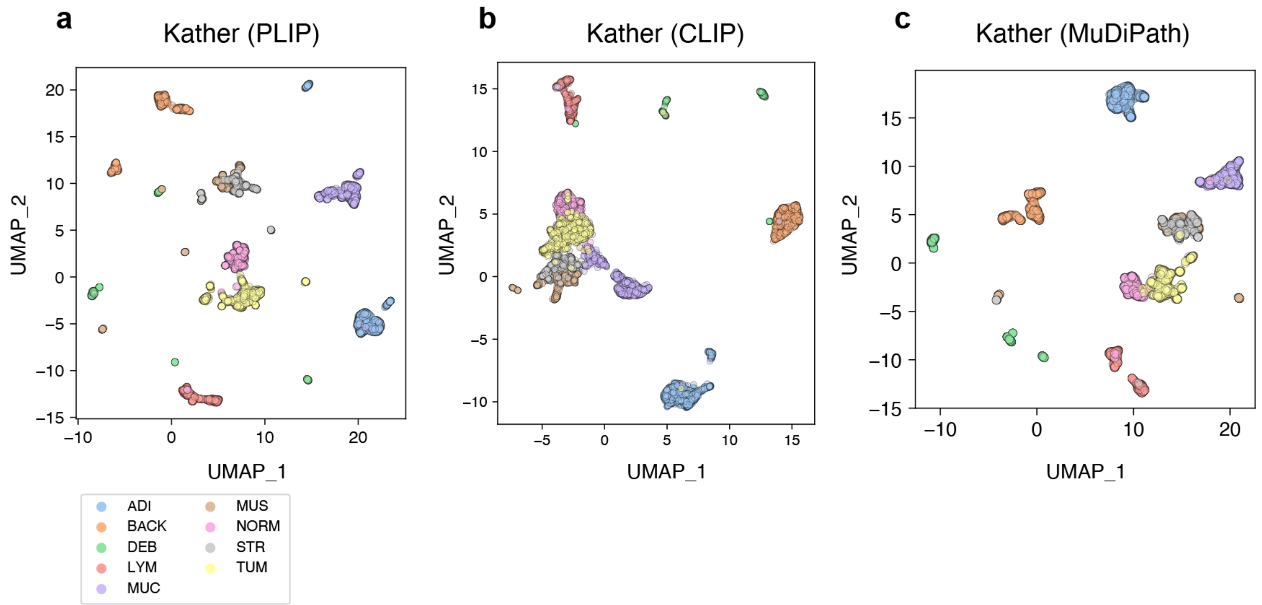


**Extended Figure 4**. Comparison of image embeddings between **(a)** PLIP, **(b)** CLIP, and **(c)** MuDiPath models in the Kather colon dataset. ADI: Adipose tissue, BACK: background, DEB: debris, LYM: lymphocytes, MUC: mucus, MUS: smooth muscle, NORM: normal colon mucosa, STR: cancer-associated stroma, TUM: colorectal adenocarcinoma epithelium.

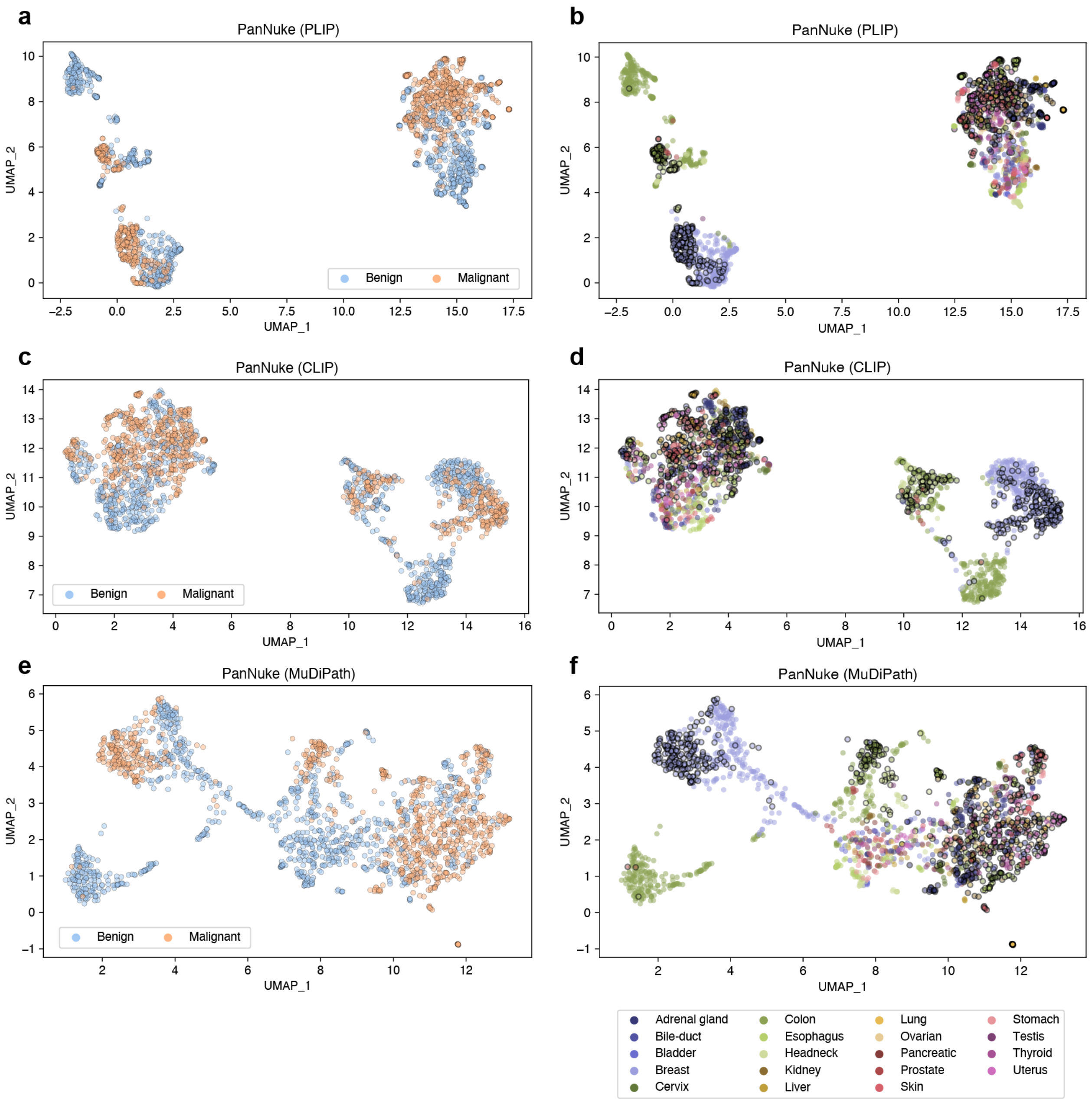


**Extended Figure 5**. Comparison of image embeddings between PLIP, CLIP, and MuDiPath models for the PanNuke dataset. (**a**) Image embeddings generated by the PLIP model, colored by benign and malignant. (**b**) Image embeddings generated by the PLIP model, colored by 19 pathology subspecialties. Scatters with black edges indicate malignant images. (**c**) Image embeddings generated by the CLIP model, colored by benign and malignant. (**d**) Image embeddings generated by the CLIP model, colored by 19 pathology subspecialties. Scatters with black edges indicate malignant images. (**e**) Image embeddings generated by the MuDiPath model, colored by benign and malignant. (**f**) Image embeddings generated by the MuDiPath model, colored by 19 pathology subspecialties. Scatters with black edges indicate malignant images.

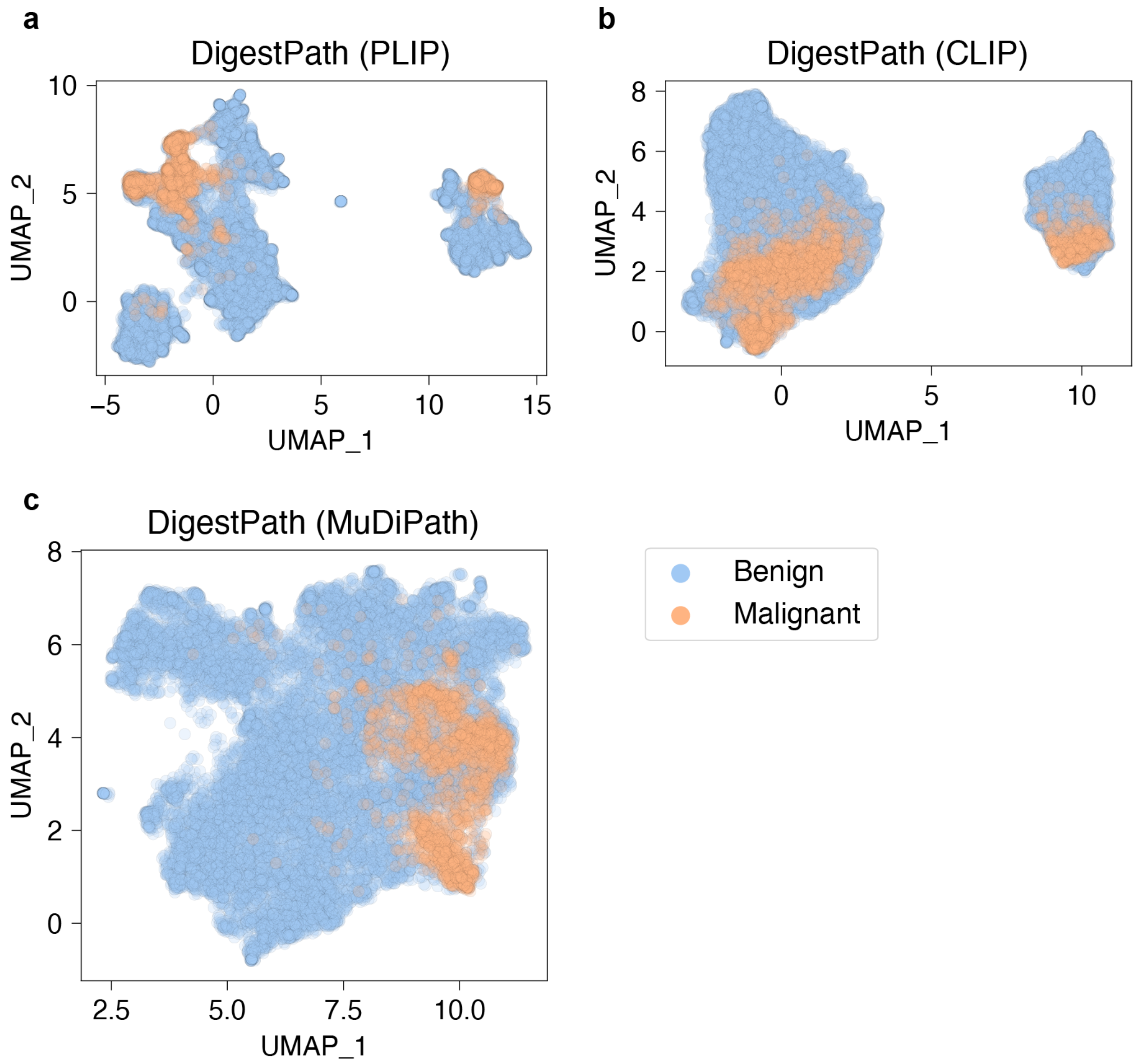


**Extended Figure 6**. Comparison of image embeddings between **(a)** PLIP, **(b)** CLIP, and **(c)** MuDiPath models in the DigestPath dataset.

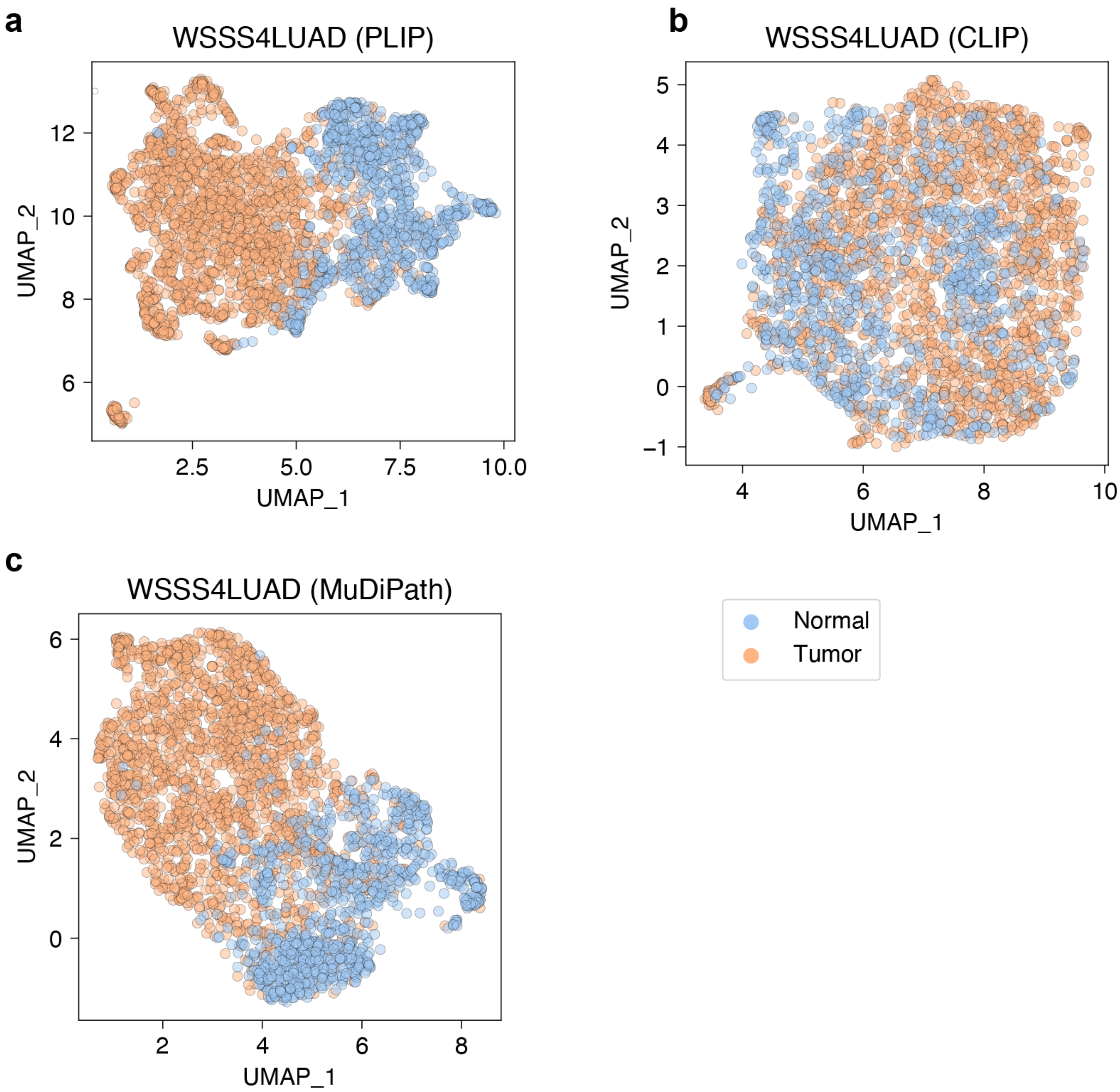


**Extended Figure 7**. Comparison of image embeddings between **(a)** PLIP, **(b)** CLIP, and **(c)** MuDiPath models in the WSSS4LUAD dataset.

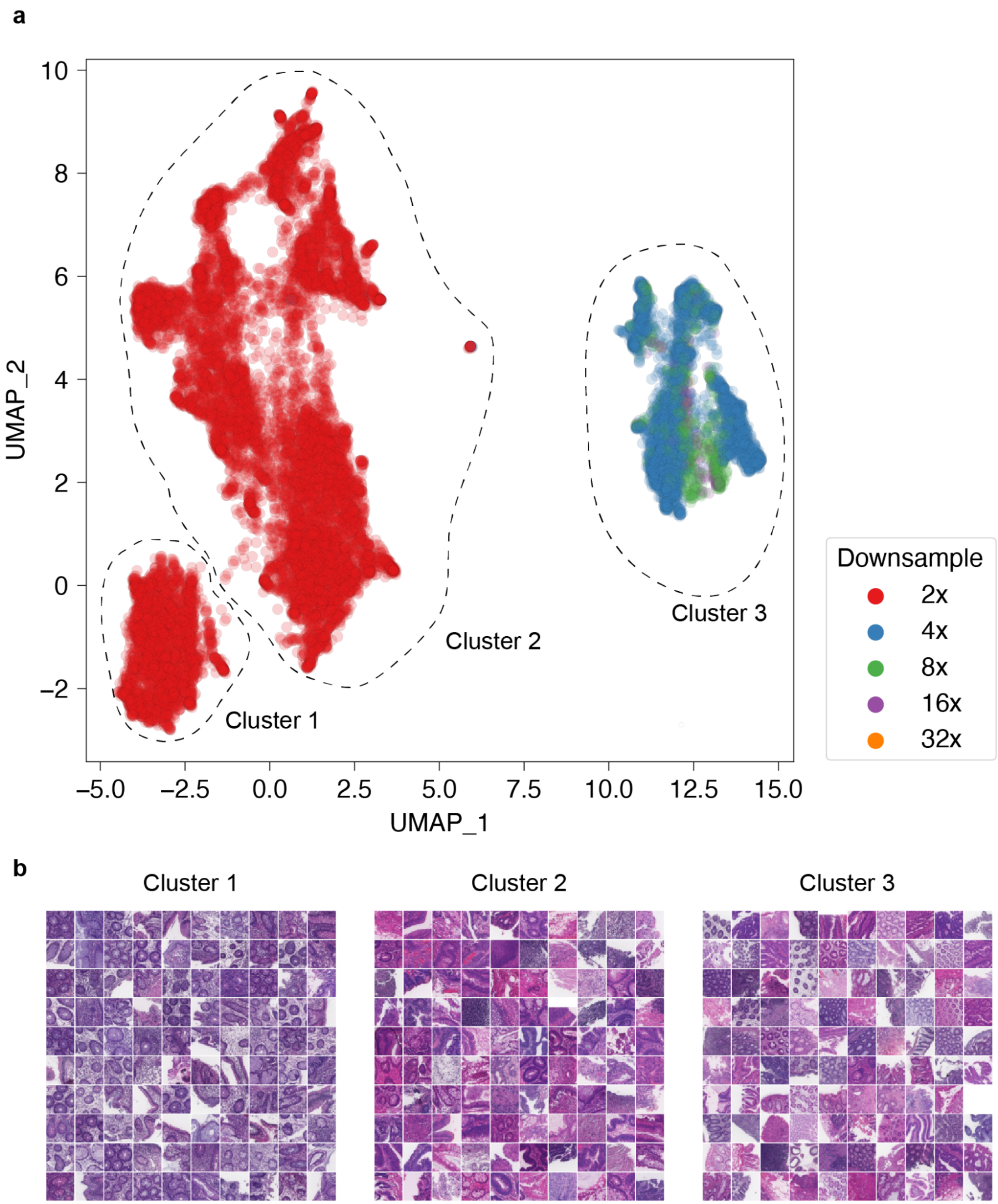


**Extended Figure 8**. Cluster visualization in DigestPath. (**a**) Image patches in low-dimensional space colored by different downsampling rates. (**b**) Visualization of image patches on different clusters.

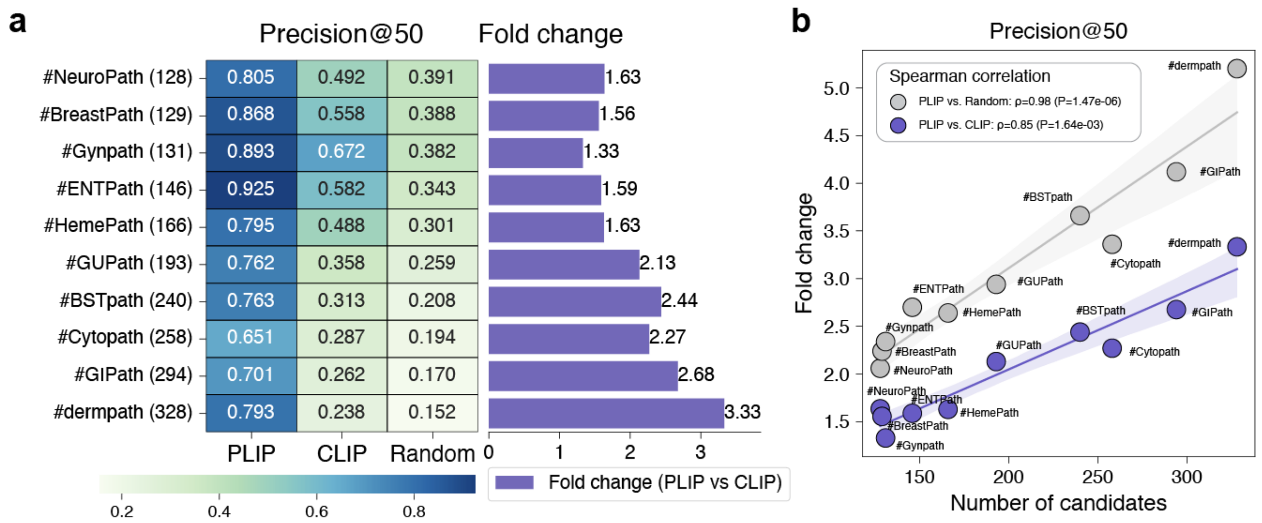


**Extended Figure 9**. Text-to-image retrieval performances for Precision@50. **(a)** Image retrieval performances for Precision@50 within each of the pathology subspecialty-specific hashtags. **(b)** Spearman correlations between the number of candidates and fold changes for Precision@50 when comparing the PLIP model with random and CLIP, respectively.

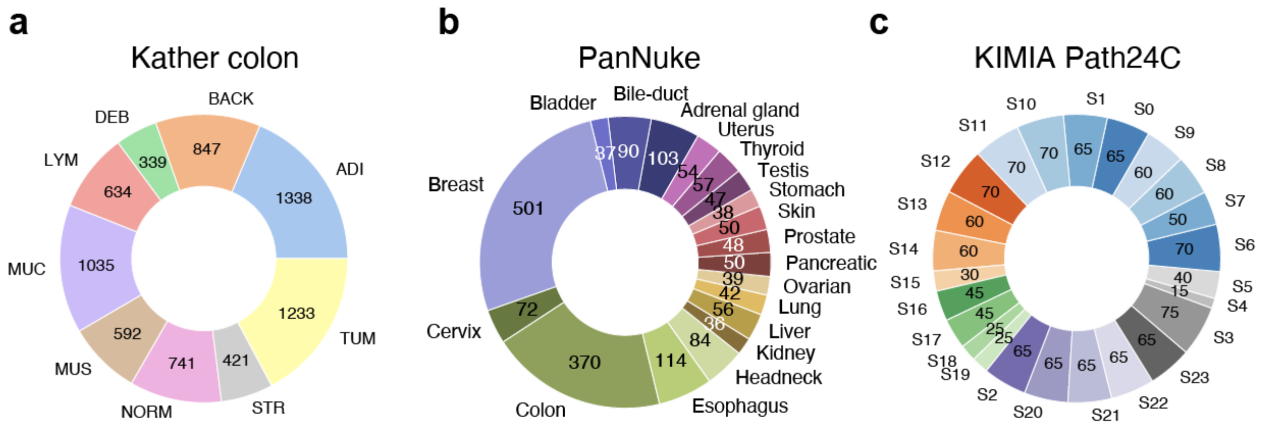


**Extended Figure 10**. External validation datasets used for image-to-image retrieval. **(a)** The Kather colon dataset (9 colon tissue types). **(b)** The PanNuke dataset (19 pathology subspecialties). **(c)** The KIMIA Path24C (24 staining textures).
